## Supplementary figure 1 for "Lack of CCDC146, a ubiquitous centriole and microtubule-associated protein, leads to non-syndromic male infertility in human and mouse"

| | | | |
|---|---|---|---|
| H.sapiens | 1 | ----MEDSSTDTEKEEEEEKD---------------------EKDQE-PI | 24 |
| P.troglodytes | 1 | ----MEDSSTDTEKEEEEEKD---------------------EKDQE-PI | 24 |
| M.mulatta | 1 | ----MDSINT---------------------------------------- | 6 |
| C.lupus | 1 | ----MEDSTTDTEQEEEEEKD---------------------EKDQEDPI | 25 |
| B.taurus | 1 | ----MEDSTTDTDQEEEEEKG---------------------GKAQEDPI | 25 |
| M.musculus | 1 | ----MEYNSKDAEEETEDEEEGLKEAEGEEQEKEEVTATSETDDDQEDLP | 46 |
| R.norvegicus | 1 | ----MEDRSKYIAEESEDEED---EEQEEKEKKGGASTSTETEEDQEDIP | 43 |
| G.gallus | 1 | MRNLQKMSQAGDESSSSDEKE--------------------LEVEKEQPI | 30 |
| D.rerio | 1 | --MMRQAQQRSSSPSHDDQQE---------------------SSERATPI | 27 |
| X.tropicalis | 1 | MSSLGEEKAESSDSSTEEEEE-----------------------VEEKPL | 27 |
| H.sapiens | 25 | YAIVPTINIQDERFVDLSETPAFIFLHELHAMGKLPGTRMAALKAKYTLL | 74 |
| P.troglodytes | 25 | YAIVPTINIQDERFVDLSETPAFIFLHELHAMGKLPGTRMAALKAKYTLL | 74 |
| M.mulatta | 7 | --------------------------LELHAMGKLPGTRMAELKAKYTLL | 30 |
| C.lupus | 26 | YAIAPTINIQDERCVDLSTTPAFICLHELHSLGKLPGTRMAELKAKYTLL | 75 |
| B.taurus | 26 | YATAPTINIQDERFVDLSTTPAFICLHELHALGKLPGTRMAELKAKYTLL | 75 |
| M.musculus | 47 | SVLVPTVNIREERLINLAETPAFLCLHELHSKGKLPGTRMAELKAKYTLL | 96 |
| R.norvegicus | 44 | SVIVPTINIREERLVDLAQTPAFLCLNELHTSGKLPGTRMAELKAKYTLL | 93 |
| G.gallus | 31 | CALAPTVYTRDEGSTDVTSSPAFQCLDELFSAGKLTSSRVAELKAKYNLL | 80 |
| D.rerio | 28 | SALAPEVDLNQEQPAVLSLNPALQCLEQLFSTGKISVEKVARLKASFNLL | 77 |
| X.tropicalis | 28 | YPVAPS-KVWGGDTKDIYASLAFQYLDELATSGKISGTRTAELKEKYTLL | 76 |
| H.sapiens | 75 | HDAVMSTQESEVQLLQNAKRFTEQIQQQQFHLQQADNFPEAFSTEVSKMR | 124 |
| P.troglodytes | 75 | HDAVMSTQESEVQLLQNAKRFTEQIQQQQFHLQQADNFPEAFSTEVSKMR | 124 |
| M.mulatta | 31 | HDAVMSTQESEVQLLQNAKRFTEQVQQQQFHLQQADNFPEAFSTEVSKMR | 80 |
| C.lupus | 76 | HDTVVSTQESEVQLLQDAKRFTEQIQQQQFHLQRADDFPEASTTEVSKMR | 125 |
| B.taurus | 76 | HDTVMSTQESEVQLLQDAKRFTEQIQEQQFHLQQADNFPQAFTTEVSKMR | 125 |
| M.musculus | 97 | HDTVVSTQESEVQLLENAKRFTEQIQQQQVCLQQAEDFPNVFTTEVCKLR | 146 |
| R.norvegicus | 94 | HDTVVSTQESEVQLLENAKRFTEQIQQQQVRLQEADDFPNVFTTEVSKLR | 143 |
| G.gallus | 81 | HETVRSLQESEIQLLQEAKRLSVDLEQQQHELEKAEQFPEESSSEVCRIR | 130 |
| D.rerio | 78 | NETLRSSQDSEMQLLLEAKTLRMTLEQQRQELENAELFPEGPDTEVSRLK | 127 |
| X.tropicalis | 77 | YQTLKRVQESEVKLLQDAKCFTNQLEQQEW--EKSDQLYEDYNTEISTLR | 124 |
| H.sapiens | 125 | EQLLKYQNEYNAVKEREFHNQYRLNSLKEEKIIIVKEFEKITKPGEMEKK | 174 |
| P.troglodytes | 125 | EQLLKYQNEYNAVKEREFHNQYRLNSLKEEKIIIVKEFEKITKPGEMEKK | 174 |
| M.mulatta | 81 | EQLLKYQNEYNAVKEREFHNQYRLNSLKEEKIIIMKEFEKIPKPGEMEKK | 130 |
| C.lupus | 126 | EQLLKYQNEYNAVKERESHTQYRLNSLTEEKNLIIKEFEKIPKPGEMEKK | 175 |
| B.taurus | 126 | EQLLKYQNEYNAVKEREFHNQYRLNSLVEEKNLILKEFEKIPKPGEMEKK | 175 |
| M.musculus | 147 | EQLLKYQNEYTAAQEREYNIQYRLTSLTEEKSIILKEFEKIPKPGEIEKK | 196 |
| R.norvegicus | 144 | EQLLKYQNEYTAVQEREYHTQYRLNSLTEEKSLVLKEFEKIPKPGEIEKK | 193 |
| G.gallus | 131 | QQLLNCQNEYNAIKEREHEIQFRIKCLQEEKRLLEKEYERIPKQGEADKK | 180 |
| D.rerio | 128 | QTLLQYYNQLKEDEERDYQMQFQLECLKEEKLYLEKEYQDQPKQAELESR | 177 |
| X.tropicalis | 125 | EQLLKYNNDINEAEEREYSLQYKLECLQEEKKLLQREYDRMPKAGDYEKK | 174 |
| H.sapiens | 175 | MKILRESTEELRKEIMQKKLEIKNLREDLASKQKQLLKEQKELEELLGHQ | 224 |
| P.troglodytes | 175 | MKILRESTEELRKEIMQKKLEIKNLREDLASKQKQLLKEQKELEELLGHQ | 224 |
| M.mulatta | 131 | MKILRESTEELRKEIMQKKLEIKNLREDLASKQKQLLKEQKELEELLEHQ | 180 |
| C.lupus | 176 | MRQLRESTEELRKEIMQRKLEIKNLREDLASKQKQLLKEQKELEELLECQ | 225 |
| B.taurus | 176 | MRLLRESTEELRKEVMQKKLEIKNLREDLTSKQKQLLKEQKELEELLEYQ | 225 |
| M.musculus | 197 | TRLLKESTEELRKEVIQRRLEIKNLREDVVLKQKQLVREQKELEELMEYQ | 246 |
| R.norvegicus | 194 | TKLLKESTEELRKEVIQKRLEIKNLREDVVSKQKQLMKEQKELEELMDYQ | 243 |
| G.gallus | 181 | MRKMKEECDELHKEVIQRKAEVNAMKEDISSKQELMLIDKKEVEKLLEDQ | 230 |
| D.rerio | 178 | YATLKNSCEEVRKEVLKLRQEVRTLTESMETQQKHIKKEQLELEDLKNRV | 227 |
| X.tropicalis | 175 | IKILQENIKELQKENTQRRMEIKALKEDLETKQRETEREQKELGQKLEQQ | 224 |
| H.sapiens | 225 | VVLK---DEVAHHQTIPVQIGKEIEKITRKKVEMEKKKIVLEQEVKTLND | 271 |
| P.troglodytes | 225 | VVLK---DEVAHHQTIPVQIGKEIEKITRKKVEMEKKKIVLEQEVKTLND | 271 |
| M.mulatta | 181 | VVLK---DEVAHHQTIPVQIGKEIEKITRKKVEMEKKKILLEQEVKNLND | 227 |
| C.lupus | 226 | VTLK---DEVVHHQAVPVQIGKEIEKTTRKKVEMEKKKIVLEYELKELND | 272 |
| B.taurus | 226 | VTLK---DEVVHHQSIPVQIGKEIEKTTRKKVEMEKKKLVLEYELKELND | 272 |

| | | | |
|---|---|---|---|
| M.musculus | 247 | VGLK---DDVVHHQSVPVQITKEIEKMTRKKVETEKKNIVLESELKELSD | 293 |
| R.norvegicus | 244 | VGLK---DDVVHHQSVPVQITKEIEKLTRRKIETEKKNVVLEFELKELSD | 290 |
| G.gallus | 231 | DSLKVTQDELVKILGVPVQLKKEAEKINQKKIDAEKKNEDLNNQMQELNS | 280 |
| D.rerio | 228 | ESNE---AELAQVLLIPVQLRKEVDRVAHKKSELEKQVAKIEQHRLDNVE | 274 |
| X.tropicalis | 225 | EVVK---HELVQLSYVPLQIGKETEKINRRIMEAQKKENMLGKEFQELND | 271 |
| H.sapiens | 272 | SLKKVENKVSAIVDEKENVIKEVEGKRALLEIKEREHNQLVKLLELAREN | 321 |
| P.troglodytes | 272 | SLKKVENKVSAIVEEKENVIKEVEGKRALLEIKEREHNQLVKLLELAREN | 321 |
| M.mulatta | 228 | SLKKVENKVSAIVEEKENVIKEVEGKRALLEIKEREHNQLVKLLELAREN | 277 |
| C.lupus | 273 | SLKKVETKINSIMEEKEDVIKEVEGKRALLEIKEREYNQLVKLLELTRVS | 322 |
| B.taurus | 273 | SLKKVETKINAVMEEKEDVIKEVEGKRALLEIKEREYNQLVKLLELTREN | 322 |
| M.musculus | 294 | SLKKLENKVNTLAEERDDIMKEVEGKRTLLEVKEREYGQLLKLLELTKEN | 343 |
| R.norvegicus | 291 | SLKKLENKVSALAEERDDTIKEVEGKRTLLEVKEREYGQLLKLLELTKEN | 340 |
| G.gallus | 281 | TLNDIDKGTEKILQEREDVMKELDGKRALLESKEQECAALARLLEIGREK | 330 |
| D.rerio | 275 | HQKSLEAKCEKVKEAIKEVVREQEGHKAQLVTAENKQDRLLTDLQMAKAK | 324 |
| X.tropicalis | 272 | NLKQMEHKAKLMLEEKEEVMKELEGKQTWLETKEQECNRLTKLLELAKEN | 321 |
| H.sapiens | 322 | EATSLTER-------------------GILDLNLRNSLIDKQNYHDELSR | 352 |
| P.troglodytes | 322 | EATSLTER-------------------GILDLNLRNSLIDKQNYHDELSR | 352 |
| M.mulatta | 278 | EATSLTER-------------------GILDLNLRNSLIDKQNYHDELSR | 308 |
| C.lupus | 323 | EATSLTER-------------------GILDLNLRNCLIDKQNYHDELSR | 353 |
| B.taurus | 323 | EASSLTER-------------------GILDLNLRNCLIDKHNYHDELSR | 353 |
| M.musculus | 344 | EASSLAER-------------------GILDINLRNCLIDKQNYHDELSR | 374 |
| R.norvegicus | 341 | EASSLAER-------------------GILDINLRNCLMDKQNYHDELSR | 371 |
| G.gallus | 331 | QLAILVER-------------------DSLNQKLNERIFEKKQQQETLIY | 361 |
| D.rerio | 325 | EAAFMDQRSLFQHFCNINAKHIQIHLVGLLEISMEQIVAEFISLRDKLAR | 374 |
| X.tropicalis | 322 | EAVALGER-------------------AAMDLSLRHAMVEKQTQHDALTR | 352 |
| H.sapiens | 353 | KQREKERDFRNLRKMELLLKVSWDALRQTQALHQRLLLEMEAIPKDDSTL | 402 |
| P.troglodytes | 353 | KQREKERDFRNLRKMELLLKVSWDALRQTQALHQRLLLEIEAIPKDDSTL | 402 |
| M.mulatta | 309 | KQREKERDFRNLRKMELLLKVSWDALTQTQAMHQRLLLEIEAIPKDDSTL | 358 |
| C.lupus | 354 | KQREKERDFRNLKKMELLLKVSWDALTQTQALHQRLLLEIEAIPKDDSTL | 403 |
| B.taurus | 354 | KQREKERDFRNLKKMELLLKVSWDALNQTQALHQRLLLEIEAIPKDDSTL | 403 |
| M.musculus | 375 | KQREKERDFRNLKKTELLLKVSLDALTQAQAMNQRLLLEMEAIPKEYLTL | 424 |
| R.norvegicus | 372 | KQREKERDFRNLKKTELLLKVSLDALSQAQALNQRLLLEMEAIPKEDLLL | 421 |
| G.gallus | 362 | MQGEKDKELRNLKKMELQMKMIHESLQQEKSQHKTLKLEAEAIPKINGVL | 411 |
| D.rerio | 375 | DTRLKDKLMLALKRNELQLKQARDALTNTQQQHERIQAQKETVSEADILL | 424 |
| X.tropicalis | 353 | KTREKERDLRNTKKMELQLKSALDALAHTQNVYEKIKAEVDCFPKADALL | 402 |
| H.sapiens | 403 | SERRRELHKEVEVAKRNLAQQKIISEMESKLVEQQLAEENKLLKEQENMK | 452 |
| P.troglodytes | 403 | SERRRELHKEVEVAKRNLAQQKIISEMESKLVEQQLAEENKLLKEQENMK | 452 |
| M.mulatta | 359 | SERRRELHKEVEVAKRNLAQQKIISEMESKLVEQQLAEENKLLKEQENMK | 408 |
| C.lupus | 404 | SERRRELHKEVEVAKRNLAQQKILSEAETKLVEQQLAEENKLLKEQENMR | 453 |
| B.taurus | 404 | SERRRELHKEVEVAKRNLTQQKILSEAESKLVEQQLAEENKLLKEQENIQ | 453 |
| M.musculus | 425 | PERRKELHKEVDLARRNLAQQKSLSEAEAKLVEQQIAQENKLLKEQESLR | 474 |
| R.norvegicus | 422 | PEQRKELHKEVDLAKRNLAQQRSLSEAEAKLVEQQIAEENKLLKEQETMR | 471 |
| G.gallus | 412 | LERRRELQKEIEMKKRCLAEQEMVSDTDARRLEECIAEEGRLFKEQEKCR | 461 |
| D.rerio | 425 | -ERRKDLEKEVENLKRNLI------------------------------- | 442 |
| X.tropicalis | 403 | -EKRKELRKEVEVIRRQFAQQQTLTEAEAHTLEQCIAEEEQLVKEQSERR | 451 |
| H.sapiens | 453 | ELVVNLLRMTQIKIDEKEQKSKDFLKAQQKYTNIVKEMKAKDLEIRIHKK | 502 |
| P.troglodytes | 453 | ELVVNLLRMTQIKIDEKEQKSKDFLKAQQKYTNIVKEMKAKDLEIRIHKK | 502 |
| M.mulatta | 409 | ELVFNLVRMTQIKIDEKEQKSKDFLKAQQKYTNIVKEIKAKDLEIRIHKK | 458 |
| C.lupus | 454 | DVVFNLVRMTQIKIDEKDQKSKDFLKAQQRYTNIVKEIKTKDLEIRIHKK | 503 |
| B.taurus | 454 | ELLFNLVRMTQIKIDEKEQKSKDFLKAQQKYTSIVREIKAKDLEIRIHRK | 503 |
| M.musculus | 475 | ELVFNLGRMTQIKIDEKEQKAKDFLKAQRRYSEIVKEIKSKDLEIRLYKK | 524 |
| R.norvegicus | 472 | EVLFNLGRMTQIKMEEKEQKAKDFLKSQRRYCDIIKEIKSKKLEIRLYRK | 521 |
| G.gallus | 462 | SELSRLAHLTWLKVEEKEQKSREVQKVQIQLQNIIKEIKRKDLEIEEYKK | 511 |
| D.rerio | 443 | --------------------------SELKYKQSKQELQGKGLVIQEHIK | 466 |
| X.tropicalis | 452 | EDLVNLTRLVQIKAEEREQKSRDLIKAKQRYQQVLQEVKGRQLIIGEHKK | 501 |
| H.sapiens | 503 | KKCEIYRRLREFAKLYDTIRNERNKFVNLLHKAHQKVNEIKERHKMSLNE | 552 |

| | | | |
|---|---|---|---|
| P.troglodytes | 503 | KKCEIYRRLREFAKLYDTIRNERNKFVNLLHRAHQKVNEIKERHKMSLNE | 552 |
| M.mulatta | 459 | KKCEIYRRLKEFAKLYDTIRNERNKFVNLLHKAHQKVNEIKERHKMSLNE | 508 |
| C.lupus | 504 | KKREIHRRLKEFAKLYDTIRNERNKFVNLLHKAHQKVNEIKERHKMLLNE | 553 |
| B.taurus | 504 | KKREIHRRLREFAKLYDTIRNERNKFINLLHKAHQKVNEIKERHKMSLNE | 553 |
| M.musculus | 525 | RKHEIHRRLREFASLYDTIRNERNKFVNLLHKAHQKVNEIKERLKMSLNE | 574 |
| R.norvegicus | 522 | RKREIHRRLKEFAGLYDAIRNERNKFVNLLHKAYQKVNEIKERLKMSLNE | 571 |
| G.gallus | 512 | RKRRVHKQLQGVANMCDVIKNERNKCMHLVNVAQQKTAEIEDRIKVKASE | 561 |
| D.rerio | 467 | QTKETQTRL----------------------------------------- | 475 |
| X.tropicalis | 502 | KNQDVQKRLKEFAKMYDIIRNERNKCVSLIQTAMQRASELREKLKIFANE | 551 |
| H.sapiens | 553 | LEILRNSAVSQERKLQNSMLKHANNVTIRESMQNDVRKIVSKLQEMKEKK | 602 |
| P.troglodytes | 553 | LEILRNSAVSQERKLQNSMLKHANNVTIRESMQNDVRKIVSKLQEMKEKK | 602 |
| M.mulatta | 509 | LEILRNSAVSQERKLQNSMLKHANNVTIRESMQNDVRKIASKLQEMKEKK | 558 |
| C.lupus | 554 | LEILRNSAVTQERKLQNSMLKHANNVTIRESMQNDVHKIVAKLQEMKEKK | 603 |
| B.taurus | 554 | LEILRNSAVTQERKLQNSMLKHANNITIRESMQNDVCKIVAKLQEMKEKK | 603 |
| M.musculus | 575 | LEILRNSAVSQERKLQNAMLKHSNNVTIKESIQNDVCKITAKLQEMKEKK | 624 |
| R.norvegicus | 572 | LEILRSSAVSQERKLQNAMLKHSNNVTIRESMQNDVCKITAKLQEMKEKK | 621 |
| G.gallus | 562 | IETLRNTLITQERELQKQHMKNKNNAAIKESLKNDCSKVAQVMYEMNEKK | 611 |
| D.rerio | 476 | --------------LQRSRLQHSHSYKLRDDLKKEVSQTLQALQEMQQNR | 511 |
| X.tropicalis | 552 | MEILRNNAVNKDRQLQKIKLKENNSQMVRDSLQKDLSRVRVYLEEMKEKR | 601 |
| H.sapiens | 603 | EAQLNNIDRLANTITMIEEEMVQLRKRYEKAVQHRNESGVQLIEREEEIC | 652 |
| P.troglodytes | 603 | EAQLNNIDRLANTITMIEEEMVQLRKRYEKAVQHRNESGVQLIEREEEIC | 652 |
| M.mulatta | 559 | EAQLNNIDRLANMITMIEEEMVQLRKRYEKAVQHRNESGVQLIEREEEVC | 608 |
| C.lupus | 604 | EAQLNNIDRLASMITTVEEEMVQLRKKYEKAVQRRNESGVQLIQREEEVC | 653 |
| B.taurus | 604 | EAQLNNIDRLANMITMIEEEMVQLRKKYERAVQRRNESGVQLIEREEEVC | 653 |
| M.musculus | 625 | EAQLTSMDRLASMITVIEEEMVQLRKKYEKAVQRRNESGVQLIEREEEVC | 674 |
| R.norvegicus | 622 | EAQLTSMDRLASMITVIEEEMVQLRKKYEKAVQRRNESGVQLIEREEEVC | 671 |
| G.gallus | 612 | KQQVLDLDGLTNAVTRIEEEIAQLHKKYKRATEEQTESGLLLRSREEEIC | 661 |
| D.rerio | 512 | EEQKLKLRKLTDTVNMLEQKKLHICKRYEAELQKRNQESVQLAEREKEWR | 561 |
| X.tropicalis | 602 | EQQKMQIGRLTNMINQAEVDMIQLRKKYQTAVQNRNERGVQLIEREEEIC | 651 |
| H.sapiens | 653 | IFYEKINIQEKMKLNGEIEIHLLEEKIQFLKMKIAEKQRQICVTQKLLPA | 702 |
| P.troglodytes | 653 | IFYEKINIQEKMKLNGEIEIHLLEEKIQFLKMKIAEKQRQIRVTQKLLPA | 702 |
| M.mulatta | 609 | IFYEKINIQEKMKLNGEIEIHLLEEKIRFLKLKIAEKQRQICVTQKLLPA | 658 |
| C.lupus | 654 | IFYEKINIQEKMKLNGEIEIHVLEEKIRFLKLKIAEKQRQIHVTQKLLPT | 703 |
| B.taurus | 654 | IFYEKINIQEKMKLNGEIELHVLEEKIRFLKLKIAEKQRQIHVTRKLLPV | 703 |
| M.musculus | 675 | IFYEKVNVQEKIKLHQDVEIHILEEKIRFLKLRIAEKQRQISVTRKLVPI | 724 |
| R.norvegicus | 672 | IFYEKMNIQDKVKLHQDIEIHILEEKIRFLKLKVAEKQRQICVTQKLVPI | 721 |
| G.gallus | 662 | ILYEKINAQEFLCRKGDIEMQATDEKISFLKMKVAEKERQIKFWLKALPM | 711 |
| D.rerio | 562 | IFNEKLGVLVKMTENSNLEMYDMEDEIRNLKMEQKEEERQNDLHKKQLSN | 611 |
| X.tropicalis | 652 | IFLEKINIQEMILRNGDLEMVSMDEKIRFLKIEAAEQKRQVEQLQKNLPN | 701 |
| H.sapiens | 703 | KRSLDADLAVLQIQFSQCTDRIKDLEKQFVKPDGENRARFLPGKDLTEKE | 752 |
| P.troglodytes | 703 | KRSLDADLAVLQIQFSQCTDRIKDLEKQFVKPDGENRARFLPGKDLTEKE | 752 |
| M.mulatta | 659 | KRSLDADLAVLQIQFSQCTDRIKDLEKQFIKPDGENRARFLPGKDLTEKE | 708 |
| C.lupus | 704 | KRALDADLAVLQIQFSQCTERIKDLEKQFINPDGENRIRFIPGKDMTQEE | 753 |
| B.taurus | 704 | KSALDADLAVLQIQFSQCTDKIKDLEKKFINPEGENRTRLLPGKDMTEEE | 753 |
| M.musculus | 725 | KKSLDADLAVIQIQFSQCTDRIKDLEKLFVNPDSKGRVRFIKGKDLTEEE | 774 |
| R.norvegicus | 722 | KKSLDANLAVVQIQFSQCADRIKALEKCFVNPDCQGRVRFIPGKDLTEEE | 771 |
| G.gallus | 712 | KRVLDAELVVLQIQYSQCKDRIKEMEEIFVDPTNESRKRDLGGKDPSAPE | 761 |
| D.rerio | 612 | KLALEEERVLLQIQLSEINDRVSELEEACVD---DNRARNLNGEDPSPEE | 658 |
| X.tropicalis | 702 | KRMLESDLVTLQIQLSQCNDRVAELETQTEDPGKEYRTRLLEGKDPSLQE | 751 |
| H.sapiens | 753 | MIQKLDKLELQLAKKEEKLLEKDFIYEQVSRLTDRLCSKTQGCKQDTLLL | 802 |
| P.troglodytes | 753 | MIQKLDKLELQLAKKEEKLLEKDFIYEQVSRLTDRLCSKTQDCKQDTLLL | 802 |
| M.mulatta | 709 | MIKKLDKLELQLAKKEEKLLEKDFIYEQVSRLTDRLCSKTQDCKQDTLLL | 758 |
| C.lupus | 754 | MIKKLDSLELQLAQKEEKLLEKDFIYEQVSRLTDRLCSKTQAYKQDTLLL | 803 |
| B.taurus | 754 | MIKKMDELELQLAKKEEKLLEKDFIYEQVSRLTDRLYSKTQACKQDTLLL | 803 |
| M.musculus | 775 | MIKKLDMLELQIAKKEEKLLEKDFIYEQVSQLTNRLKGKTQACKKDTLLL | 824 |
| R.norvegicus | 772 | MIKKLDMLELQLAKKEEKLLEKDFIYEQVSQLTNRLKTKTQACKMDTLLL | 821 |
| G.gallus | 762 | LQKKIKQLEVELVQKEEKLLETDIIYQHIARLTDRIRATAENGKQGTLLL | 811 |
| D.rerio | 659 | LIEKIEQLEIHLADKEAQLLEMELVCEQVGRLSKRIQIKADNSNEDTLHL | 708 |

| | | | |
|---|---|---|---|
| X.tropicalis | 752 | LMKKTEELELHLAQKEEQLLEKDLLYEQVSHLTKKIHTKAENGKQDTLLL | 801 |
| H.sapiens | 803 | AKKMNGYQRRIKNATEKMMALVAELSMKQALTIELQKEVREKEDFIFTCN | 852 |
| P.troglodytes | 803 | AKKMNGYQRRIKNATEKMMALVAELSMKQALTIELQKEVREKEDFIFTCN | 852 |
| M.mulatta | 759 | AKKMNGYQRRIKNATEKMMAVVAELSMKQALTIELQKEVREKEDFIFTCN | 808 |
| C.lupus | 804 | AKKMNGYRKKIKDATEQMMALVAELSMKQALAIELQKEVREKEDFIFSCN | 853 |
| B.taurus | 804 | AKKMNGYQKKIKDATEKMMALVAELSMKQAMAIELQKEVKEKEEFIFTCN | 853 |
| M.musculus | 825 | AKKMNSYQQKIKVVTQEMMALVAELSMKQALTIELQKEVREKEEFIFSCS | 874 |
| R.norvegicus | 822 | AKKMNSYQKKIKDVTQEMMALVAELSMKQALTIELQKEVREKEEFIFSCS | 871 |
| G.gallus | 812 | ATRINELQKKIKDRTQKMMALVAELSMKQALAIKLQQEMRDREEFLMIVS | 861 |
| D.rerio | 709 | AKKVNELQAQIRKRSRKMMAVVAELSMRKAECMGLQQEMKEKELQLDLCQ | 758 |
| X.tropicalis | 802 | SKKMNELHKKIKNTTQKMMSLIAELSMQQANSIKLQEELRNKEKMVEACH | 851 |
| H.sapiens | 853 | SRIEKGLPLNKEIEKEWLKVLRDEEMHALAIAEKSQEFLEADNRQLPNGV | 902 |
| P.troglodytes | 853 | SRIEKGLPLNKEIEKEWLKVLRDEEMHALAIAEKSQEFLEADNRQLPNGV | 902 |
| M.mulatta | 809 | SRIEKGLPLNKEIEKEWLKVLRDEEMHALALAEKPQSSLEADNRQMPNGV | 858 |
| C.lupus | 854 | SRIEKGLPLNKDIETEWLKILRDEEMYALAVAEKSREFLVSDNRQLPNGV | 903 |
| B.taurus | 854 | SRMEKGLPLNKEIEREWLKVLRDEEMYAMAIAEKSREFLEAETRQLPNGV | 903 |
| M.musculus | 875 | ARIEKGLPLNREIEKDWLKVLRDEEMYAFATAEMSREYMETDYRQLPNGV | 924 |
| R.norvegicus | 872 | SRIEKGLPLNREIEKDWLKVLRDEEMYAFATAEKTREYIDTDYRQLPNGV | 921 |
| G.gallus | 862 | SRIDQGLPPPKETEIEWLKVLRNEKMHKEAAEARARQAAEEEQAAVPGHV | 911 |
| D.rerio | 759 | KHVEQGLPPSDSIENEWLRHLQDQHRRLADAERKAMLAEEDEWNQLPNGV | 808 |
| X.tropicalis | 852 | TRMEQGLPPSEGTENEWKKMIRDKHRLQKEKEDKARMAEEEEQHWLPNGA | 901 |
| H.sapiens | 903 | YTTAEQRPNAYIPEADATLPLPKPYGALAPFKPSEPGANMRHIRKPVIKP | 952 |
| P.troglodytes | 903 | YTTAEQRPNAYIPEADATLPLPKPYGALAPFKPSEPGANMRHIRKPVIKP | 952 |
| M.mulatta | 859 | YTTAEQRPNAYIPEAEATLPLPKPYGALAPFKPSEPGANMRHIRKPIIKP | 908 |
| C.lupus | 904 | YTTAEPRPNAYIPEAEAALPLPKPYGALAPFKPSEQGANMRHIRKPVPKP | 953 |
| B.taurus | 904 | YTTAEQRPNAYIPEAEATLPLPKPYGALAPFKPSEPGANIRHIRKPVIKP | 953 |
| M.musculus | 925 | YTTAEQRPNAYMPEADTELPLPKPYGAMAPFKPSEPGANRRHIRKPVVKP | 974 |
| R.norvegicus | 922 | YTTAEQRPNAYIPETDATLPLPKPYGALAPFKPSEPGANRRHIRKPIIKP | 971 |
| G.gallus | 912 | LTTAEPRPTAYVPDDAYSLPVPRPYGALAPFKPSEPGSNIRHFRKPIIKP | 961 |
| D.rerio | 809 | YTTAELRPNAYIPIDD-PLPVPKHYGALAPFKPTEPGANIRHIRKPKIKP | 857 |
| X.tropicalis | 902 | YTTAEQRPNAYIPDDESALPLPRPYGILAPFKPSETGSNMRHIRKPIIKP | 951 |
| H.sapiens | 953 | VEI | 955 |
| P.troglodytes | 953 | VEI | 955 |
| M.mulatta | 909 | IEI | 911 |
| C.lupus | 954 | IEI | 956 |
| B.taurus | 954 | IEI | 956 |
| M.musculus | 975 | IEI | 977 |
| R.norvegicus | 972 | IEI | 974 |
| G.gallus | 962 | IEI | 964 |
| D.rerio | 858 | IEI | 860 |
| X.tropicalis | 952 | IEI | 954 |
