## Supplementary material for "Lack of CCDC146, a ubiquitous centriole and microtubule-associated protein, leads to non-syndromic male infertility in human and mouse": Table supplementary 1

List of primers used for Sanger verification of the identified variants by WES.

| Primer name | Primer sequence (5'-3') | Tm (°C) | Product length (bp) |
|---|---|---|---|
| CCDC146-Int8F | ATTGCTGGGTCAAACGGTAG | 57.3 | 422 |
| CCDC146-Int9R | GGCAGCAAAAACAACTTCCT | 55.3 | |
| CCDC146-Ex15F | CAGCTGATAGAGCGGGAAGA | 59.3 | 457 |
| CCDC146-Int15R | TCCAAGAAAAGCAGAAAATGC | 54.0 | |

List of primers used for knock-out mice genotyping.

| Primer name | Primer sequence (5'-3') | Tm (°C) | Product length (bp) |
|---|---|---|---|
| Ccdc146-Ex2F3 | GGGAGGAACAGGAGAAGGAG | 61.4 | 151 |
| Ccdc146-Ex2R3 | TCATGCAGACAGAGGAAAGC | 57.3 | |

List of primers used for knock-in mice (HA-Tag) genotyping.

| Primer name | Primer sequence (5'-3') | Tm (°C) | Product length (bp) |
|---|---|---|---|
| shCCD_ki-F1 | ACTTGGTGGGTGTTGTCCTA | 58.49 | 149 |
| shCCD_ki-R1 | TCCCTCCTCTTCATCTTCAGT | 57.52 | |
| lgCCD_ki-F1 | AGAAATCAGGGAGGGGTTGC | 59.67 | 504 |
| lgCCD_ki-R1 | AATTGATGAGCCGCTCCTCC | 60.18 | |
